# Tolerogenic Nanoparticles Impacting B and T Lymphocyte Responses Delay Autoimmune Arthritis in K/BxN Mice

**DOI:** 10.1101/2020.07.02.185140

**Authors:** Amrita Srivastava, Britni M. Arlian, Lijuan Pang, Takashi K. Kishimoto, James C. Paulson

## Abstract

Current treatments for unwanted antibody responses largely rely on immunosuppressive drugs compromising overall immunity. New approaches to achieve antigen-specific tolerance are desirable to avoid unwanted side effects. Several nanoparticle-based approaches designed to tolerize the B or T cell arms of the humoral immune response have shown promise for induction of antigen-specific tolerance, raising the possibility that they could work synergistically if combined. Earlier we showed that Siglec-engaging tolerance-inducing antigenic liposomes (STALs) that display both an antigen (Ag) and glycan ligands of the inhibitory co-receptor CD22 (CD22L) lead to robust antigen-specific B cell tolerance to protein antigens in naïve mice. In another approach, administration of free Ag with poly(lactic co-glycolic acid)-rapamycin nanoparticles (PLGA-R) induced robust antigen-specific tolerance through production of regulatory T cells. Here we illustrate that co-administration of STALs together with PLGA-R to naïve mice induced more robust tolerance to multiple antigen challenges than either nanoparticle alone. Moreover, in K/BxN mice that develop spontaneous autoimmune arthritis to the self-antigen glucose-6-phosphate-isomerase (GPI), co-delivery of GPI-LP-CD22L and PLGA-R delayed onset of disease, and in some mice prevented the disease indefinitely. The results show synergy between B cell-tolerizing STALs and T cell-tolerizing PLGA-R and the potential to induce tolerance in early stage autoimmune disease.

## 1. Introduction

Undesired immune responses are responsible for numerous human maladies including autoimmune diseases, ^[1–2]^ allergies, ^[3]^ transplantation rejection, ^[4–5]^ and production of inhibitory antibodies to biotherapeutic medicines. ^[6]^ Existing treatments for these conditions are not antigen-specific and therefore lead to broad immune suppression with numerous adverse effects. ^[7]^ Thus, to suppress undesired immune activation, antigen-specific tolerance approaches are needed that provide immune tolerance to the antigen of interest while leaving the rest of the immune system intact.

Various approaches have been used to induce antigen-specific immune tolerance in the B cell and T cell arms of the humoral immune response. ^[8–11]^ A direct approach to induce antigen-specific tolerance in B cells invokes the recruitment of inhibitory coreceptors from the sialic acid-binding immunoglobulin-like lectin (Siglec) family to the B cell receptor (BCR), in particular the B cell Siglecs CD22 and Siglec-G/10. ^[12–14]^ Co-presentation of antigen with ligands of CD22 or Siglec-G on liposomal nanoparticles results in targeting of B cells that recognize the antigen and recruitment of the Siglecs to the immunological synapse formed when a B cell receptor (BCR) engages the antigen. ^[15–17]^ This engagement leads to a potent apoptotic signal resulting in elimination of antigen-reactive B cells, leaving the rest of the B cell repertoire unaffected, and induction of B cell tolerance due to depletion of the antigen-specific B cells. ^[18]^ The utility of **S**iglec **t**olerizing **a**ntigenic **l**iposomes (STALs) has been successfully demonstrated in several models of disease, one involving the generation of inhibitory antibodies to FVIII in hemophilia mice, ^[15]^ another involving tolerization of animals to allergen sensitization in a peanut antigen anaphylaxis model, ^[19–20]^ and a third involving apoptosis of autoantigen-specific memory B cells from RA patients. ^[21]^ STALs are particularly effective in naïve animals since there is minimal activation of CD4+ T cells that are required for B cells to differentiate into cells that produce high affinity antibodies of the IgG class, and are less suitable for inducing tolerance in animals with an ongoing immune response with antigen specific CD4+ T helper or T memory cells.

An alternative for inducing antigen-specific tolerance is inducing tolerance in the T cell arm of the immune response. Currently available to allergic patients is allergen immunotherapy (AIT) where gradually increasing doses of antigen are administered by injection or orally to patients over several years to build up tolerance. While AIT has been helpful in desensitizing patients to antigen, there is limited long lasting desensitization when treatment ends. ^[20, 22–23]^ Another approach aimed at treating ongoing immune responses involves targeting of antigen experienced T cells with peptide-MHC bound nanoparticles to induce T regulatory cells (Tregs), but this technology requires knowledge of the precise peptide antigen recognized by the T cells. ^[24–25]^ Other approaches seek to program T cells through antigen-presenting cells (APCs) presented with antigen coupled to syngeneic cells, ^[26–27]^ or synthetic particles carrying antigens and/or immune-modulators. ^[28–35]^ The general strategy is to create tolerogenic APCs that secrete cytokines to induce antigen-specific anergy/deletion of T cells, or reprogram them to Tregs. ^[10–11, 24]^

A particularly attractive option currently in clinical development involves administering of antigen with poly(lactic co-glycolic acid) (PLGA) particles containing the immunomodulator rapamycin. Initially, Maldonado *et al*. showed that PLGA nanoparticles encapsulated with both antigen and the immunomodulatory drug rapamycin could induce robust tolerance to a variety of antigens in naïve animals. ^[29]^ Subsequently, it was shown that PLGA nanoparticles with rapamycin (PLGA-R) co-injected with soluble antigen produced equally robust tolerance. ^[30–31, 34–36]^ Mechanistic studies support the idea that the non-targeted rapamycin-PLGA particles efficiently deliver the drug to APCs to create a tolerogenic cytokine environment, which in turn induces Tregs that prevent activation of T helper cells needed by B cells to produce inhibitory antibodies. ^[29–31,36]^

Because the tolerogenic nanoparticle technologies impact immune responses of the B cell and T cell arms of the immune system by different mechanisms, we reasoned that they might be synergistic, providing more robust tolerance than either alone. Initial attempts to incorporate rapamycin into the lipid bilayer of STALs showed improved tolerance induction in naïve mice, but were limited by the dose of rapamycin that could be achieved and had insignificant impact on tolerance induction in previously sensitized mice. ^[37]^ Here we have compared the two tolerogenic nanoparticle platforms alone or in combination under various conditions of antigen presentation and tested them for preventing inflammatory disease in the antigen-specific K/BxN arthritis model. Administering STALs and PLGA-R in combination significantly suppressed antibody production to liposomal-antigen over either nanoparticle alone. In K/BxN mice that develop autoimmune rheumatoid arthritis (RA) dependent on production of antibodies to the self-protein, glucose-6-phosphate isomerase (GPI), ^[38–40]^ neither GPI-STALs or PLGA-R alone had significant impact on the onset of disease. When administered together weekly for up to 5 weeks, the onset of disease was delayed, and some mice remained disease free for up to 150 days. Some benefit could also be achieved by delaying treatment until ankle swelling was apparent. The results suggest that combining these B and T cell tolerance strategies hold promise for suppressing unwanted immune reactions.

## 2. RESULTS AND DISCUSSION

### 2.1. Nanoparticle synthesis and characterization

Unless otherwise noted, STALs and PLGA-R nanoparticles were prepared as previously described with few modifications. ^[15, 29]^ For incorporation into liposomes (LP), the mouse CD22 ligand (CD22L) was coupled to pegylated-lipid (**Figure 1A**) and the protein antigen (Ag) was also coupled to lipid as described. ^[15]^ Schematic illustrations of STALs (Ag-LP-CD22L) and PLGA-R nanoparticles are shown in Figure 1B.

**Figure 1.**
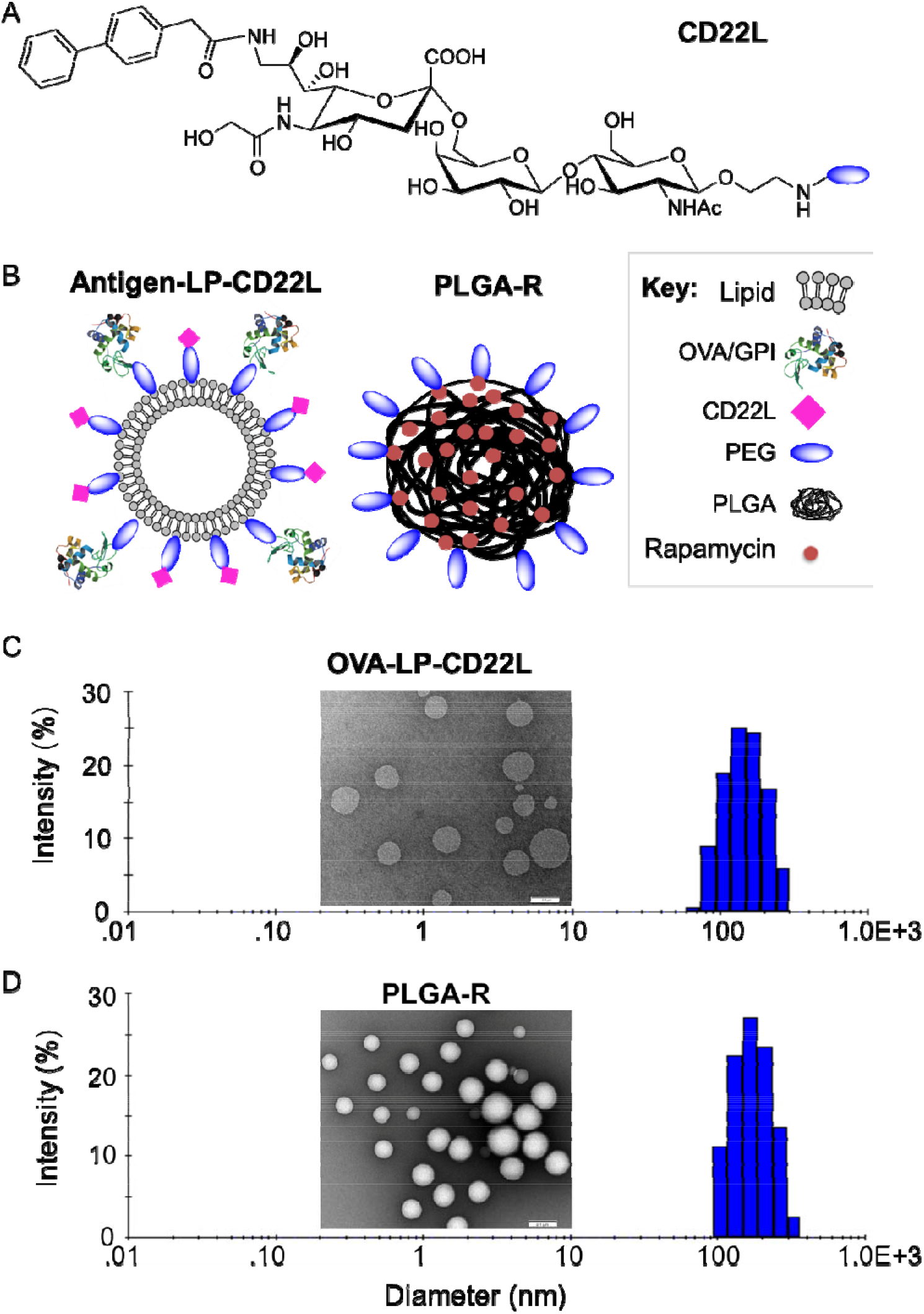
Nanoparticle formulation and characterizations. (A) Chemical structure of the murine CD22 ligand linked to PEG. (B) Schematic illustration of targeted OVA/GPI-STALs (OVA/GPI-LP-CD22L) and PLGA rapamycin (PLGA-R) nanoparticles. (C-D) Representative transmission electron microscopic images of OVA-LP-CD22L (inset C) and PLGA-R (inset D), and hydrodynamic size distribution of OVA-LP-CD22L (156±32; plot C) and PLGA-R (182±31; plot D) as determined by dynamic light scattering.

STALs with ovalbumin (OVA) as antigen (OVA-STALs; OVA-LP-CD22L) and biodegradable PLGA-R nanoparticles were characterized by dynamic light scattering (DLS) and transmission electron microscopy (TEM) (Figure 1C-D). The hydrodynamic diameter of OVA-LP-CD22L and PLGA-R was found to be 156±32 and 182±31, respectively, as measured by DLS. TEM characterization confirmed that both OVA-LP-CD22L and biodegradable PLGA-R showed uniform spherical morphology.

### 2.2. STALs and PLGA-R synergize to produce enhanced tolerance

Since STALs are formulated with protein antigen, STALs and PLGA-R were tested alone and in combination for their ability to suppress antibody production against liposomal ovalbumin (OVA-LP or OVA-LP-CD22L) in C57BL/6J mice. The concentration of OVA (0.1 mol %) and CD22 ligand (1.5 mol %) used in the STALs was selected based on previously optimized formulations. ^[15, 37]^ PLGA-R nanoparticles contained 50-100 μg of rapamycin as earlier studies indicated this as an optimal dose for inducing tolerance. ^[34–35]^ Similar formulations of STALs and PLGA nanoparticles have been reported to have serum half-lives in mouse blood of 11 and 16 hours, respectively. ^[41–42]^ Mice were treated (i.v.) once (day 0), or twice (days 0 and 14) with OVA-LP, OVA-STALs (OVA-LP-CD22L) or OVA-LP and PLGA-R, followed by challenge i.p. 14 and 35 days later with OVA/Alum and free OVA (fOVA), respectively. With one treatment, analysis of anti-OVA serum titers showed that treatment with OVA-STALs or PLGA-R alone or together significantly reduced serum titers of OVA relative to mice treated with OVA-LP. Moreover, OVA-LP-CD22L and PLGA-R together further reduced anti-OVA IgG1 titers relative to mice treated with STALs or OVA-LP + PLGA-R alone, demonstrating synergy between the two tolerogenic nanoparticle systems (**Figure 2A**).

**Figure 2.**
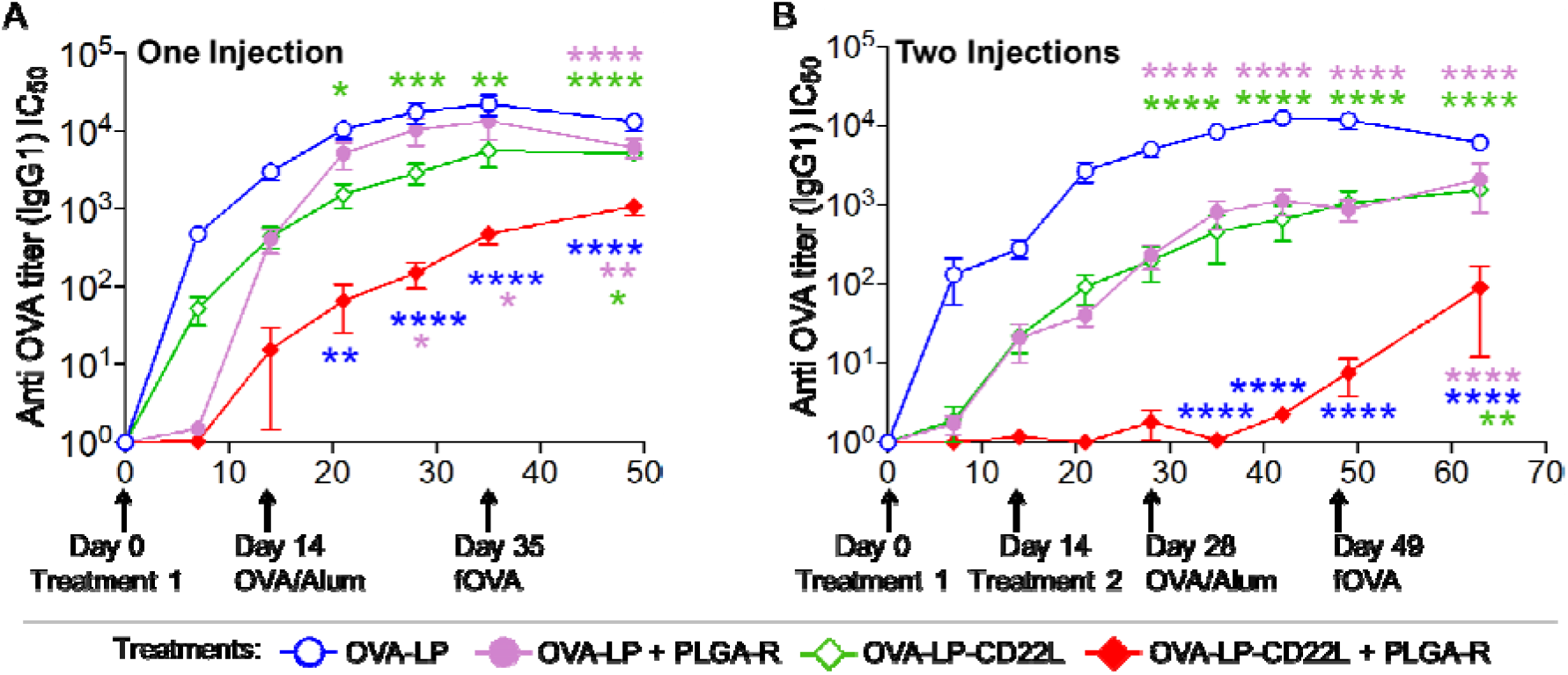
Co-delivery of OVA-STALs (OVA-LP-CD22L) and PLGA-R nanoparticles induce greater tolerance to liposomal OVA than either nanoparticle alone. (**A**) C57BL/6J mice (*n* = 16) were immunized on days 0 with the indicated nanoparticle treatments i.v. and then challenged i.p. with OVA/Alum on day 14 and fOVA on day 35. Data were pooled from three independent experiments (**B**) C57BL/6J mice (*n* = 5) were immunized on days 0 and 14 with the indicated nanoparticle treatments i.v. and then challenged i.p. with OVA/Alum on day 28 and fOVA on day 49 and results are representative of two independent experiments. PLGA-R nanoparticles contained 100 μg of rapamycin ^[35]^. Mice were bled weekly after the treatment, and IgG1 titers were assessed for OVA. Data were normalized by dividing the titers by naïve mean and are shown here. All data represent the mean ± SEM. All statistical analyses were performed on raw data using two-way ANOVA with Tukey’s post-test (**** *P* ≤ 0.0001; *** *P* ≤ 0.001; ** *P* ≤ 0.01; \**P* ≤ 0.05).

With two treatments at 0 and 14 days, greater tolerance to OVA challenge was evident with the combined treatment with OVA-LP-CD22L + PLGA-R showing significantly greater suppression than either nanoparticle alone (**Figure 2B**). Notably, while OVA-LP + PLGA-R induced weak suppression of anti-OVA production, two treatments with fOVA + PLGA-R induced strong tolerance equivalent to OVA-STALs + PLGA-R (see online supplementary figure S1). Thus, we suggest that PLGA-R is more effective at inducing tolerance with monovalent antigen than the multivalent antigen exemplified by OVA-LP.

Since the strategy for combining STALs and PLGA-R envisioned the induction of CD4+ antigen specific regulatory T cells that would synergize with STALs for suppression of B cell activation ^[29]^, we investigated production of Tregs in naïve mice. OVA specific CD4+ T cells (OTII cells) were adoptively transferred into naïve mice and the next day mice were treated with OVA-STALs (OVA-LP-CD22L) alone or with one or two doses of OVA-STALs + PLGA-R. On day 28 post-treatment, the number of Foxp3+CD25+OTII T cells in spleen were evaluated (see online supplementary figure S2). While no Treg induction was observed with OVA-STALs alone, Tregs were induced when OVA-STALs were administered together with PLGA-R nanoparticles.

### 2.3. Treatment of K/BxN arthritis mice with tolerogenic nanoparticles delays onset of disease symptoms

To evaluate the potential for the combined STALs + PLGA-R nanoparticle treatments to modulate autoimmune disease, we chose the K/BxN autoimmune rheumatoid arthritis (RA) model. ^[38–40]^ The K/BxN arthritis model recapitulates many histological features of human rheumatoid arthritis, including cartilage and bone destruction, leukocyte invasion, synovitis, and pannus formation. This is an ideal model since it is caused by development of autoimmune antibodies to the self-antigen glucose-6-phosphate isomerase (GPI), and both B and T cells are required for disease onset and pathology. K/BxN mice express the TCR transgene KRN and MHC class II molecule A^g7^, spontaneously produce GPI reactive antibodies, and develop chronic RA symptoms as they age. ^[38–40]^ Since the relevant antigen for the K/BxN model is GPI, an abundant self-antigen made by every cell, K/BxN mice are not “naïve” to the antigen, even at birth. All mice develop symptoms of RA and begin to exhibit ankle swelling at day 25-30 after weaning, allowing treatments to begin prior to disease onset. For these experiments, GPI-STALs were prepared in a manner analogous to OVA-STALs. ^[15]^

K/BxN mice were injected i.v. with 1, 2, or 5 doses of nanoparticles every 6-7 days, starting at 21-25 days of age, and monitored for disease progression by measuring front and hind paw thickness three times per week (**Figure 3A**). Mice were considered to be at the study endpoint when one or more joint(s) measured 4 mm. GPI-STALs (GPI-LP-CD22L) and PLGA-R nanoparticles administered separately in two doses failed to alter disease progression since the median survival of mice (percentage of mice without disease) treated with either nanoparticles was not significantly different from untreated mice (37, 39, and 36 days for untreated, GPI-LP-CD22L-treated, and PLGA-R-treated mice, respectively; Figure 3B). However, when GPI-LP-CD22L and PLGA-R were co-delivered to K/BxN mice using the 1, 2, or 5-time dosing scheme (Figure 3C), disease progression was significantly delayed, demonstrating synergy of combining the two nanoparticles. The delay in disease progression was dose-dependent, as the median survival of mice without disease was 47.5, 56, and 80 days for mice treated with 1, 2, or 5 doses of GPI-LP-CD22L + PLGA-R, respectively. Astonishingly, none of the mice treated 5 times with GPI-LP-CD22L+ PLGA-R had disease by 70 days of age, twenty days after dosing ended, and a few mice remained disease free to the end of the 150 day study.

**Figure 3.**
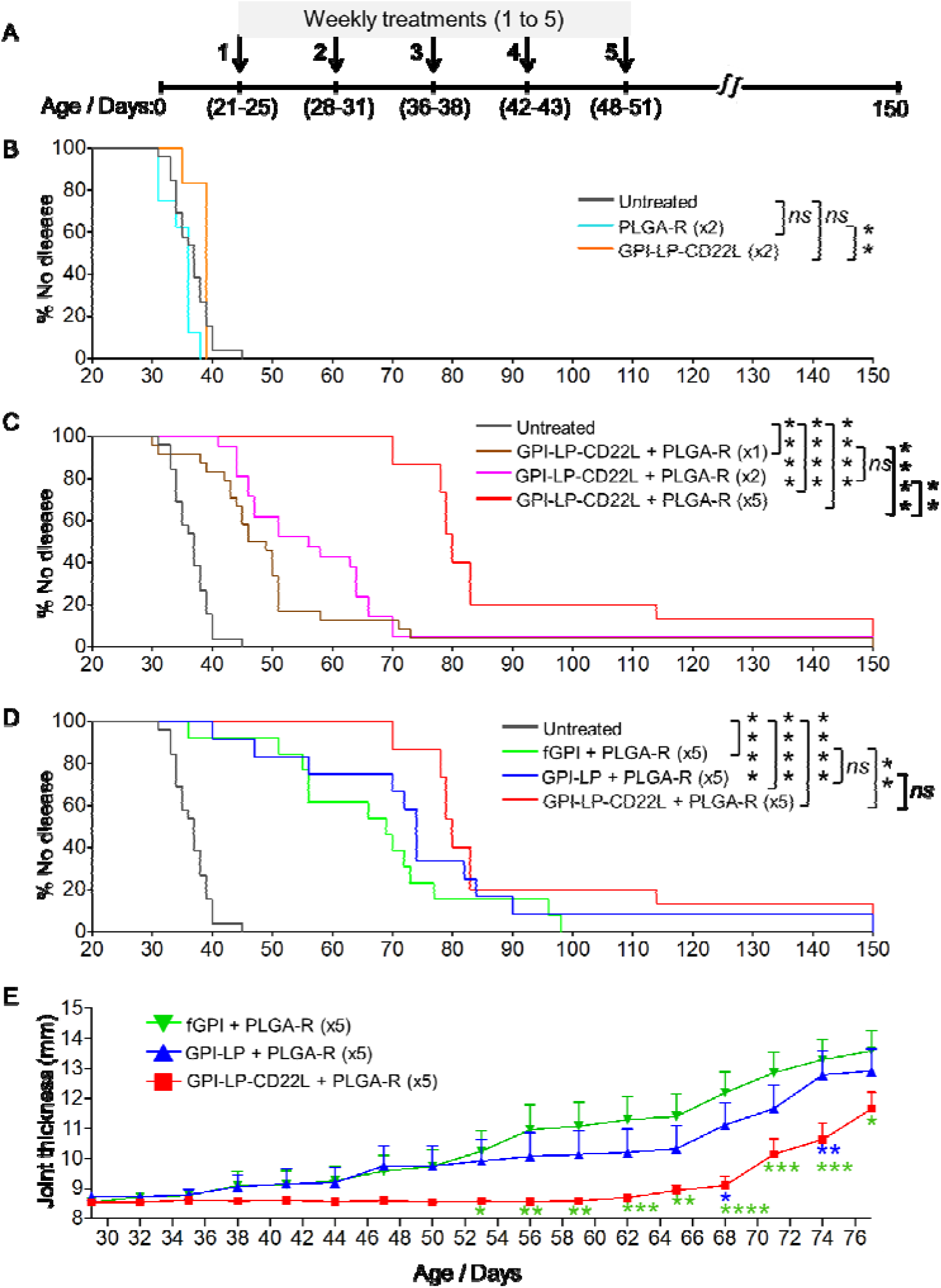
Administration of nanoparticles to K/BxN mice prior to onset of arthritis delayed disease. (**A**) Strategy for treating K/BxN mice with one, two, or five doses of nanoparticles prophylactically. PLGA-R nanoparticles contained 100 μg of rapamycin. (**B-D**) K/BxN mice, 21-25 days of age, were untreated or treated once (x1) or weekly (6-7 days) for 2 (x2) or 5 (x5) consecutive weeks. Joint measurements were performed three times a week, and mice were considered to be at the study endpoint when one or more joint(s) measured 4 mm. Results over time for each group (*n* = 6-26), are shown as percent of mice with no joint swelling over 4 mm. Results are representative of two experiments. To facilitate comparisons, results for untreated mice and GPI-LP-CD22L (GPI-STALs) + PLGA-R are shown in panels **B-D** and **C-D**, respectively. (**E**) Disease severity assessed as total joint thickness for all four paws in treated mice out to 77 days. Statistical analyses were performed using log-rank test in figure **B-D** and by two-way ANOVA with Tukey’s post-test in **E** (**** *P* ≤ 0.0001; *** *P* ≤ 0.001; ** *P* ≤ 0.01; \**P* ≤ 0.05 and ns indicates not significant).

Although we assumed that mice had abundant endogenous free antigen, we also investigated treatment of mice with PLGA-R with exogenous free soluble GPI (fGPI) or GPI liposomes (GPI-LP) without CD22L. Accordingly, mice were injected with fGPI (25 μg) + PLGA-R or GPI-LP + PLGA-R using the 5-dose injection scheme. Co-injection of PLGA-R with exogenous fGPI or GPI-LP dramatically delayed onset of disease (Figure 3D). However, in both of the antigen + PLGA-R groups, 25-40% of the mice developed disease before the end of dosing at 50 days. This delay in onset of disease can be seen in mean disease severity for each group, measured as the sum of ankle thickness for all four paws over time (Figure 3E). Thus, while antigen + PLGA-R (Figure 3D) was clearly better than PLGA-R alone (Figure 3B), treatment with GPI-STALs + PLGA-R provided a further significant delay (Figure 3D, 3E), and was the only treatment that delayed the onset of disease for all mice during treatment and for nearly 20 days after treatment ended.

To assess disease progression histologically, ankle sections from untreated wild-type C57BL/6J mice (**Figure 4A-D**) and K/BxN mice at 55 days that were untreated (Figure 4E-H) or treated with five weekly doses of GPI-LP-CD22L+PLGA-R (Figure 4I-L) were stained with Safranine O/Fast Green to assess protection of the joints under the different treatment conditions. Normal cartilage characterized by maintenance of SafO staining, chondrocyte numbers, and the bone matrix was seen in the treated K/BxN mice and untreated wild-type mice. In addition, the synovium in treated K/BxN mice and untreated wild-type mice was healthy, without the severe inflammation and infiltration of immune cells into the joint space that was evident in the untreated K/BxN mice. Finally, minimal bone resorption at the cartilage/bone margin was seen in the treated K/BxN mice and untreated wild-type mice compared to severe bone resorption in the untreated K/BxN mice. Together these histochemical data suggest treatment with the nanoparticles can prevent damage to the cartilage, bone, and joint space.

**Figure 4.**
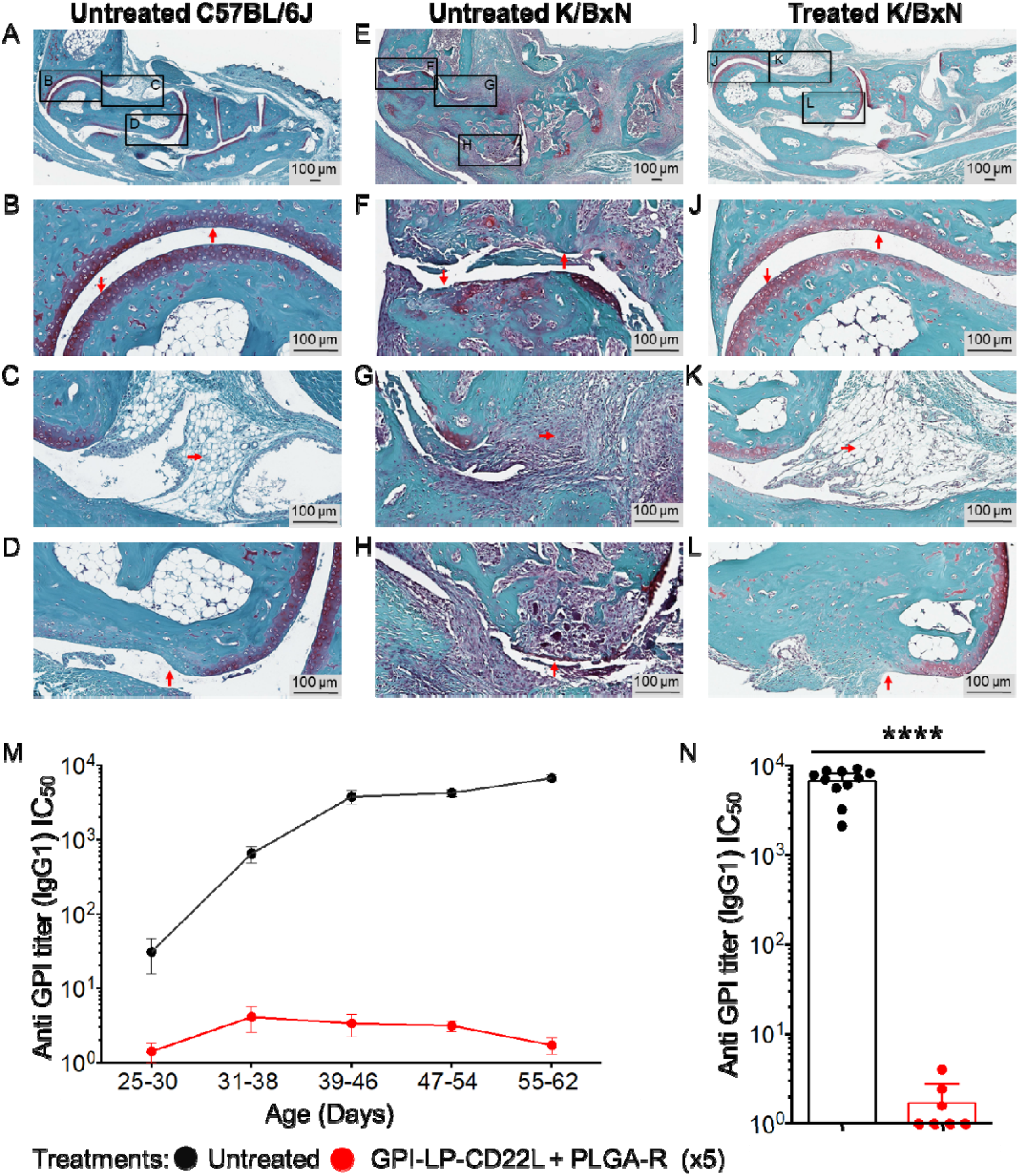
Co-administration of GPI-LP-CD22L and PLGA-R protects K/BxN mice from joint damage. Histochemical staining of day 55 paraffin-embedded ankle sections with Safranine O/Fast Green and hematoxylin counterstain reveals gross differences in the joint health of control wild-type mice left untreated (**A**), K/BxN diseased mice left untreated (**E**), and K/BxN diseased mice treated 5 times with GPI-LP-CD22L and PLGA-R (**I**). Higher magnification reveals distinct maintenance of cartilage (**J**), preservation of synovial tissue (**K**), and minimal bone resorption (**L**) in treated K/BxN mice, comparable to healthy controls (**B, C**, and **D**), which is in stark contrast to the loss of cartilage (**F**), severe infiltration of immune cells into the synovium (**G**), and severe bone resorption (**H**) seen in diseased K/BxN mice left untreated. Images are representative of two mice for each condition. (**M**) Co-administration of GPI-LP-CD22L and PLGA-R to K/BxN mice suppresses production of anti-GPI serum titers. Anti-GPI IgG titers from untreated K/BxN mice (black, *n* = 11) versus mice co-administered GPI-LP-CD22L and PLGA-R (red, *n* = 7) for 5 consecutive weeks as determined by ELISA. (**N**) Anti-GPI IgG1 titers plotted for days 55-62. All data represent the mean ±SEM. Statistical analysis was performed using the Mann-Whitney test (**** *P* ≤ 0.0001).

We also assessed the serum anti-GPI IgG titers from treated and untreated K/BxN mice by ELISA (Figure 4 M-N). While untreated K/BxN mice developed high anti-GPI titers, mice treated with GPI-STALs + PLGA-R for 5 consecutive weeks, maintained very low anti-GPI titers, providing direct evidence that the treatments suppressed antibody production to GPI.

### 2.4. Therapeutic treatment of K/BxN arthritis mice delays progression of disease

Although K/BxN mice are born with the propensity to develop autoimmune joint disease mediated by antibodies to GPI, we considered that once the disease symptoms had started, progression may be resistant to treatment with GPI-STALs + PLGA-R. However, motivated by the delayed onset of arthritis when treatment was initiated soon after weaning, we sought to test for efficacy when the first treatment was initiated after disease onset when at least one paw was swollen to a thickness of 2.8-3.05 mm. Mice were then either left untreated or given 5 weekly doses of GPI-LP-CD22L + PLGA-R containing either 50 or 100 μg of rapamycin (**Figure 5A**). For this study, we used total joint thickness, the sum of the thickness of all four paws, as a measure of disease severity. For untreated mice (*n* = 17), disease progressed rapidly with all mice achieving a score of >12 mm by day 40. Of the mice administered with GPI-LP-CD22L + PLGA-R containing 50 μg of rapamycin per treatment (Figure 5B, *middle panel, n = 17*), all but one mouse showed a delay in achieving a score of 12, and 9 of 17 mice showed a more significant delay. Remarkably, 3 mice either had no progression, or progressed and then reversed disease severity. Moreover, 2 out of 5 mice administered with GPI-LP-CD22L + PLGA-R containing 100 μg of rapamycin per treatment showed no progression (Figure 5B, right panel). These results suggest that the combination of GPI-STALs + PLGA-R is able to significantly delay or prevent disease in a prophylactic modality and even reverse the disease progression in a therapeutic treatment modality. We further analyzed the anti-GPI titers for the mice left untreated versus mice treated with GPI-STALs + PLGA-R on days 30-35 and 50-55 (Figure 5C). While anti-GPI titers for the mice left untreated or treated with 50 μg/dose increased significantly over this period, there was no significant increase in titers for mice treated with 100 μg/dose rapamycin.

**Figure 5.**
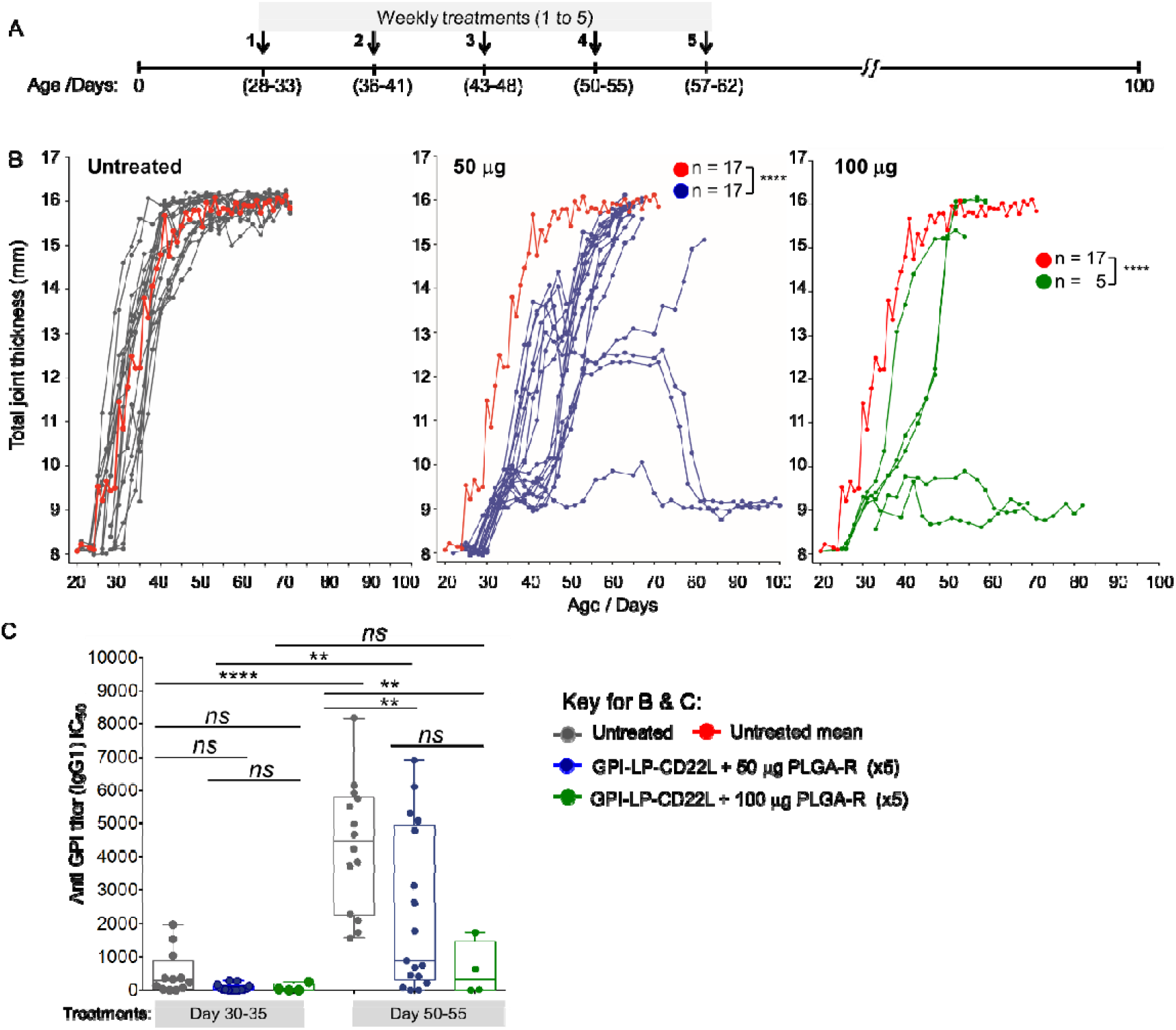
Treatment of K/BxN mice after disease onset reduces disease progression. (A) K/BxN mice were treated after disease onset measured as >3mm in one or more joints. Mice were either left untreated or treated with 5 weekly injections of both GPI-STALs (GPI-LP-CD22L) and PLGA-R nanoparticles. (B) Progression of disease severity measured by total joint thickness for untreated mice (*left panel*) and mice treated with GPI-STALs (GPI-LP-CD22L) and PLGA-R nanoparticles containing either 50 μg (*middle panel*) or 100 μg (*right panel*) of rapamycin. The mean progression for untreated mice (*red curve left panel*) is repeated in the *middle* and *right* panels for comparison. (C) Anti-GPI antibody titers from untreated and GPI-STALs + PLGA-R treated K/BxN mice on day 30-35 and day 50-55. Statistical analyses were performed using mixed-effect analysis with a Sidak post-test in **5B** and One-way ANOVA followed by Tukey’s test in **5C**(**** *P* ≤ 0.0001, ** *P* ≤ 0.01 and *ns* indicates not significant).

## 3. Conclusions

We found that co-administration of STALs targeting B cells that recognize GPI and PLGA-R that induces antigen specific T cell tolerance act synergistically to induce more robust antigen-specific tolerance compared to either nanoparticle alone. Repeated dosing was beneficial, which was particularly evident in the delay of disease onset and progression in the K/BxN arthritis model. Co-administration of 5 doses of GPI-LP-CD22L and PLGA-R prior to display of arthritis delayed disease onset until 11 weeks, with some mice remaining disease-free for the duration of the 150-day study. Therapeutic treatment of K/BxN mice with GPI-LP-CD22L and PLGA-R, after disease onset, when anti-GPI IgG1 titers had increased, showed prevention and, in some cases, even reversal of disease. Together results suggest the potential of the combined treatments to treat early stage autoimmune diseases, particularly in view of the fact that STALs have been demonstrated to suppress antigen specific human memory B cells from rheumatoid arthritis patients, ^[21]^ and that B cell depleting therapy such as rituximab (anti-CD20) is therapeutically effective in a rheumatoid arthritis setting. ^[43–44]^

## 4. Experimental Section

### Sugar-lipid conjugation

Reaction of murine CD22 ligand (9-biphenylacetyl-*N*-glycolylneuraminic acid-α-2-6-galactose-β-1-4-*N*-acetylglucosamine-β-ethylamine (6’-^BPA^NeuGc)) with NHS-PEG2000-DSPE (NOF America) gave sugar-lipid conjugate (Figure 1A) as reported previously.^[41]^ 1.5 Mol % of mCD22 ligand conjugated lipid was used for the Ag-LP-CD22L formulation.

### Protein-lipid conjugation

Ovalbumin (OVA; Worthington Biochemical LS003062) or purified recombinant mouse glucose-6-phosphate isomerase (GPI) expressed in *E. coli* (pGEX-4T-3-gpi expression vector kindly gifted by Diane Mathis and Christophe Benoist (Harvard Medical School)) was conjugated to PEGylated distearoylphosphoethanolamine (PEG2000-DSPE) using maleimide chemistry.^[45]^ Proteins were first reacted to approximately 2.5 equivalents of sulfosuccinimidyl 6-[3’-(2-pyridyldithio)propionamido]hexanoate (sulfo-LC-SPDP, Pierce), a heterobifunctional crosslinker, to introduce a thiol group. The reaction mixture was shaken at RT for 1 h, purified on Sephadex G-50, and treated with 1,4-dithiothreitol (25 mM) at RT for 10 min. Extent of protein modification was calculated by quantification of released thiol 2-pyridyl by absorbance at 343 nm. After completion of reaction, thiol-derivatized protein so obtained was purified on Sephadex G-50 column and reacted with maleimide-PEG2000-DSPE (10 eq, NOF America) under nitrogen overnight at RT. Lipid-modified protein was purified by passage over a Sephadex G-100 column and stored at 4 °C. Under these reaction conditions, proteins were modified with 1-3 lipids as determined by increased apparent MW by SDS-PAGE (see online supplementary figure S3). The final concentration of protein-lipid conjugate was calculated based on the absorbance at 280 nm.

### Liposome formulation

Liposomes were prepared by the thin film hydration method as reported previously^[15]^ using distearoyl phosphocholine (DSPC) (Avanti Polar Lipids), cholesterol (Sigma-Aldrich), and PEGylated lipids in a molar ratio of 57:38:5. The total molar fraction of PEGylated lipids, consisting of PEG2000-DSPE (Avanti Polar Lipids), ^BPA^NeuGc-PEG2000-DSPE, and protein-PEG2000-DSPE, was kept constant at 5%. DSPC, cholesterol and PEG2000-DSPE were dissolved in chloroform. After mixing appropriate amount of all the lipids, the chloroform was removed under a stream of nitrogen gas. To the dried lipid thin film, 1.5 mol % of ^BPA^NeuGc-PEG2000-DSPE (DMSO stock) was added, and this mixture was lyophilized overnight. The dried lipids were then hydrated with 0.1 mol % of protein(OVA/GPI)-PEG2000-DSPE^[37]^ in PBS to yield a lipid concentration of 5 mM. This hydrated mixture was sonicated for at least 5×30 sec with 10 min intervals. Liposomes were passed a minimum of 20 times through 800-nm, 200-nm, and 100-nm polycarbonate membranes (Avanti Polar Lipids) using a mini-extrusion device (Avanti Polar Lipids). The liposomes were purified over a CL-4B column and detected using a Nanodrop2000 (Thermo Scientific) at 280 nm. The prepared liposomes were stored at 2.5 mM concentration, based on molar content of total lipids, in PBS at 4 °C. Hydrodynamic diameter of the liposomes was found to be 156±32 nm as determined by dynamic light scattering (DynaPro NanoStar). Concentrations of liposomes are based on lipid content, and the amounts of CD22L and protein antigen as mol % based on total lipids.

### PLGA nanoparticle formulation

PLGA nanoparticles were prepared using the oil-in-water single emulsion method as described previously with slight modification.^[29]^ PLGA, a 75:25 D,L-lactide/glycolide ratio and mPEG-PLGA with 50:50 D,L-lactide/glycolide ratio were dissolved in 1mL of dichloromethane in 3:1 ratio. To this solution, 13% w/w rapamycin was added. PLGA drug solution was added dropwise to an aqueous PVA solution while vortexing, followed by sonication under ice. The single emulsion so obtained was added to 30mL of phosphate buffer, stirred for 2.5 h at RT to evaporate dichloromethane. The subsequent NP suspension was washed twice by centrifuging at 27000 rpm at 4 °C and then resuspended in 1X DPBS. The hydrodynamic diameter was found to be around 182 ±31 nm based on dynamic light scattering. PLGA NPs injections contain 50 or 100 μg of rapamycin content.

### Drug loading and encapsulation efficiency

To measure drug loading of rapamycin, PLGA-R stock (20 μL) was mixed with water (130 μL). An equal amount of acetonitrile was added to this solution to make 50/50 acetonitrile/water medium and then the mixture was sonicated for 30 min and vortexed time to time. Prior to HPLC analysis, the mixture was filtered using a 0.45 μm PVDF syringe filter. The resulting free drug was assayed using an Agilent 1100 HPLC series with a quaternary pump equipped with an Alltech C18, 5 μm, 4.6 × 150 mm column. Rapamycin absorbance was measured by a UV-vis detector at 278 nm and a retention time of 13 min in 1 mL/min using 50/50 acetonitrile/water mobile phase. Peak area of each sample was calculated using ChemStation software (Agilent). Standard solutions of known rapamycin concentrations from 30-210 μg/mL were used to make calibration curve. Rapamycin drug loading was 10-11% by weight. Rapamycin encapsulation efficiency (EE) was found to be 83% as calculated using the following equation.

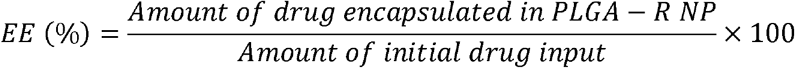

### Animals

Wild-type C57BL/6J mice were obtained from rodent breeding colony of The Scripps Research Institute (TSRI). All experimental procedures involving mice in this work were approved by the Institutional Animal Care and Use Committee of The Scripps Research Institute (La Jolla, CA).

### Immunization and blood collection

Liposomal and PLGA-R nanoparticles were delivered via the lateral tail vein in a volume of 200 μL comprising 150 μM of liposome based on total lipid concentration containing 0.1 mol % of protein (OVA/GPI) and 1.5 mol% of CD22 ligand and 50 or 100 μg of rapamycin containing PLGA-R.^[34–35]^ For OVA challenges, mice were injected intraperitoneally with 100 μg of OVA (Worthington Biochemical LS003062) plus Alum (Thermo Scientific, 77161) or 100 μg of free OVA. Serum for ELISA was obtained by centrifuging (15,000 *g*, 1 min) whole blood (50 μL, retro-orbital bleed).

### ELISAs

The levels of Anti-OVA and anti-GPI IgG titers were measured by ELISA. Assay microplates were coated with protein (50 μL/well, 100 μg/ml in PBS) and incubated overnight at 4 °C. Plates were washed with TBS-T (4 times, 0.1 % Tween 20) and blocked with 1% BSA at RT for 1 h. Serial dilutions of serum sample were added and plates were incubated (50 μL/well) at RT for 1 h. Plates were then washed 4 times and incubated with anti-IgG1 mouse-horseradish peroxidase (HRP) conjugate (1:2000 dilution; Santa Cruz Biotechnology Inc.) at RT for 1 h. After 1 h, plates were washed and developed in 75 μL/well HRP at RT for 15 min and quenched with 75 μL/well 2N H_2_SO4. Absorbance at 450 nm was measured using a Synergy H1 microplate reader (BioTek). Anti-IgG1 titers were calculated with Prism (GraphPad Software) by applying a standard four-parameter IC_50_ function.

### Statistics

Statistical significance was determined using repeated measures one- or two-way analysis of variance (ANOVA) with Tukey post-test for multiple comparisons, mixed-effect analysis followed by Sidak test, Mann-Whitney test, or the log-rank Mantel-Cox test using the Graphpad Prism 8.0. *P* <0.05 was considered significant.

### K/BxN arthritis model

K/BxN mice on a C57BL/6J background, which express both the T cell receptor (TCR) transgene KRN and the MHC class II molecule A^g7^, spontaneously produce autoantibodies recognizing glucose-6-phosphate isomerase (GPI) and develop inflammatory arthritis. For prophylactic treatment, K/BxN mice were injected i.v. with 200 μL of nanoparticles every 6-7 days for 1, 2 or 5 weeks, starting at 22-25 days of age. Nanoparticles injected include PLGA-R containing 100 μg of rapamycin mixed with 1) liposomes decorated with 0.1 mol % recombinant glucose-6-phosphate isomerase (GPI) protein and 1.5 mol % BPA CD22 ligand (GPI-LP-CD22L), 2) liposomes decorated with 0.1 mol % GPI (GPI-LP), or 3) 25 μg of free GPI (fGPI). For therapeutic treatment, K/BxN mice were left untreated or injected i.v. with 200 μL of GPI-LP-CD22L and PLGA-R containing 50 or 100 μg of rapamycin every 6-7 days, starting when the thickness of at least one paw measured 2.8-3.05 mm. Disease progression was monitored by paw thickness (front and hind) measurements collected 3 times per week using digital Vernier calipers and approximation of GPI-specific IgG titers by weekly serum ELISAs. K/BxN mice were produced through breeding of KRN and A^g7^ mice provided by the late Dr. Kerri Mowen (The Scripps Research Institute) with permission from Dr. Diane Mathis and Dr. Christophe Benoist (Harvard Medical School).

### Histology

Mouse ankles were fixed for 24 hours in buffered zinc formalin (Anatech Ltd), decalcified for 9 days in Shandon TBD-2, dehydrated by passage through increasing concentrations of EtOH, transferred to xylene, and paraffin-embedded. Tissue sections (3 μm) were mounted on glass slides, deparaffinized using Pro-Par Clearant (Anatech Ltd), and rehydrated in successive baths of absolute EtOH, decreasing concentrations of EtOH, and finally distilled water. Slides were stained in filtered hematoxylin (Ricca Chemical Company 353532) for 4 min and then washed in distilled water, acid alcohol (0.5% HCl in 70% EtOH), distilled water, Scott’s Tap Water (0.2% w/v sodium bicarbonate, 2% w/v magnesium sulfate), and finally distilled water. Slides were then immersed in 0.2% Fast Green FCF (MP Biomedicals 211922) for 6 min, washed in 1% glacial acetic acid, and stained with 0.003% Safranine O (Acros Organics 146640250) for 4 min. After staining, tissues were subsequently dehydrated as described above and cover slipped using Refrax mounting medium (Anatech Ltd). Slides were scanned with a Leica SCN400 slide scanner, and images were captured using SlidePath’s Digital Image Hub. Specimens were analyzed primarily for abnormalities in cellularity, Safranine O stain distribution, and surface fibrillation.

## Supporting Information

Supporting Information is available from the Wiley Online Library or from the author.

## Acknowledgements

We thank the following individuals at The Scripps Research Institute: Kevin Worrell for synthesis of the mCD22 ligand, Merissa Olmer for the histochemical staining protocol, Dr. Theresa Fassel for microscopy, and Anna Tran-Crie for help with preparation of the manuscript. This work was funded in part by NIH grants R01AI099141 & R01AI050143.

Received: ((will be filled in by the editorial staff))

Revised: ((will be filled in by the editorial staff))

Published online: ((will be filled in by the editorial staff))

## Table of contents

This study shows that B cells targeted STALs and T cells tolerizing PLGA-R nanoparticles work synergistically to induce profound tolerance to subsequent antigen challenges and delay onset of disease in the K/BxN mouse model of rheumatoid arthritis and in some mice prevented disease indefinitely. These two nanoparticle platforms work synergistically represents an important advance for developing treatments for autoimmune diseases and undesired immune responses.

**Keyword:** Immune tolerance

Amrita Srivastava, Britni M. Arlian, Lijuan Pang, Takashi K. Kishimoto and James C. Paulson^*^

**Tolerogenic Nanoparticles Impacting B and T Lymphocyte Responses Delay Autoimmune Arthritis in K/BxN Mice**

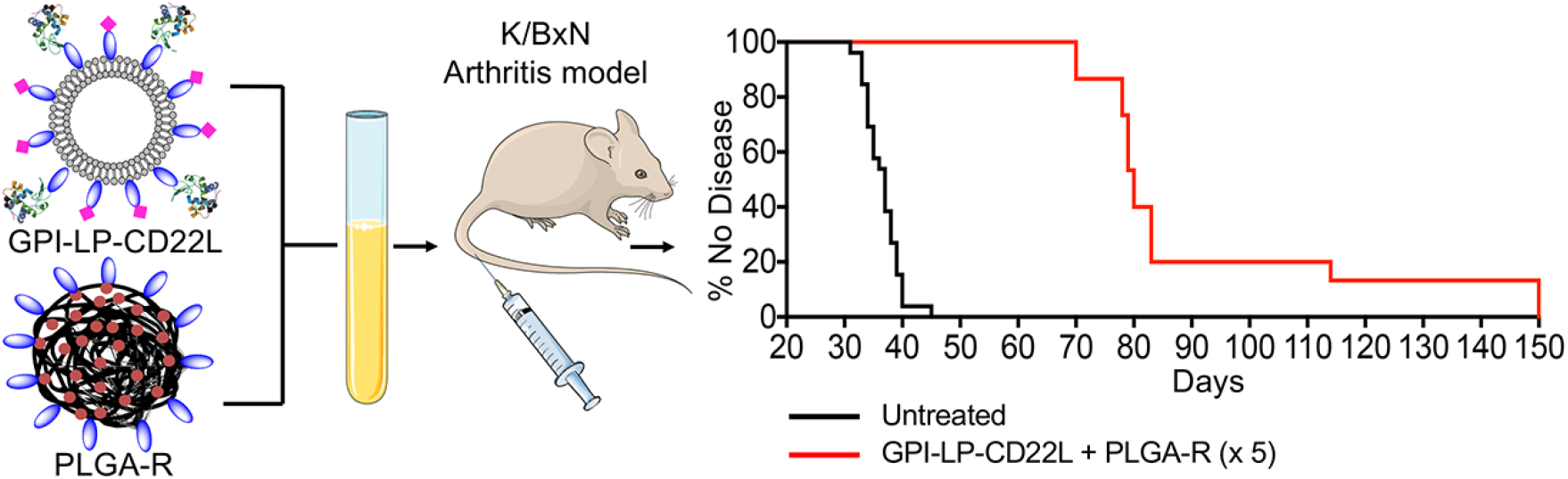

