## Supporting Information for "Tolerogenic Nanoparticles Impacting B and T Lymphocyte Responses Delay Autoimmune Arthritis in K/BxN Mice"

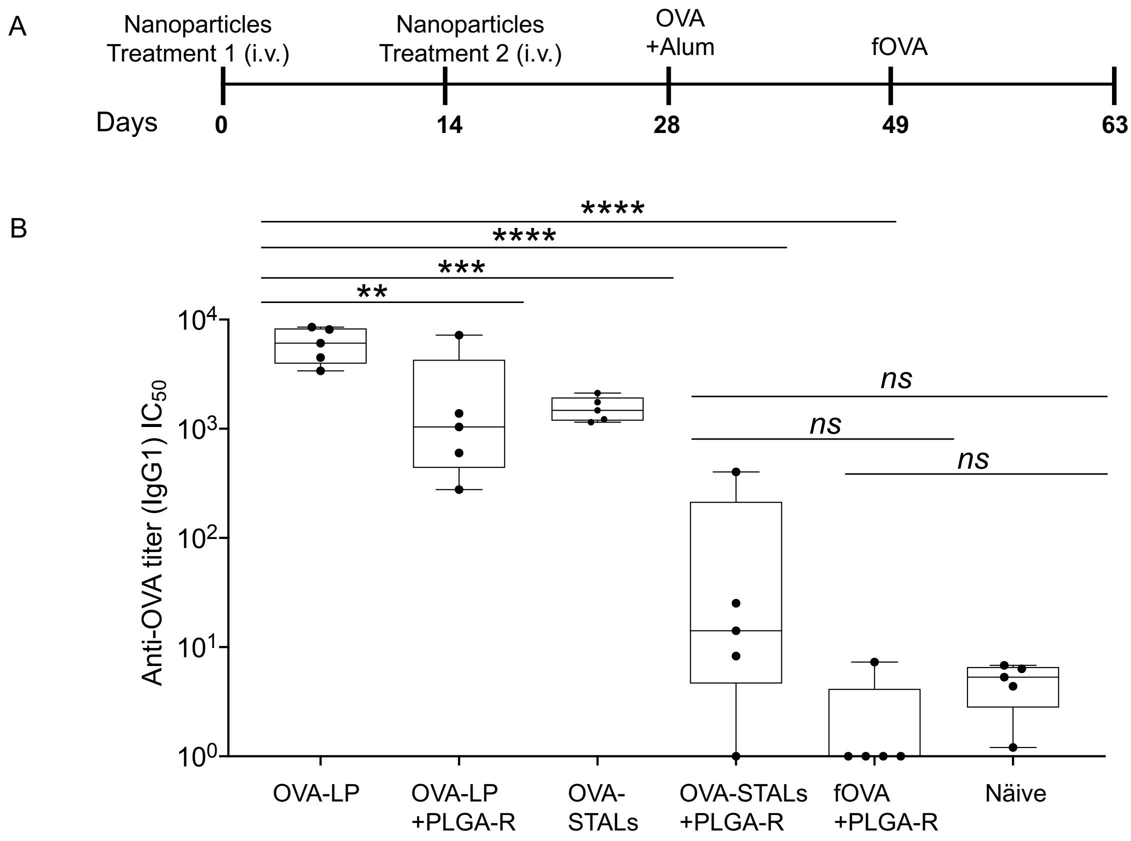


**Figure S1.** Anti-OVA titers following two treatments with PLGA-R with OVA-LP, OVA-STALs or free OVA. Shown are results summarizing the experiment described in Figure 2B with the addition of a group treated with free OVA (fOVA) and PLGA-R, comparing final anti-OVA titers at day 63. (A) C57BL/6J mice ($n$ = 5) were immunized on days 0 and 14 with the two doses of indicated nanoparticle treatments i.v. and then challenged i.p. with OVA/Alum on day 28 and fOVA on day 49 and results are representative of two independent experiments. PLGA-R nanoparticles contained 100 µg of rapamycin. Mice were bled weekly after the treatment, and IgG1 titers are shown in figure 2B for complete time course. (B) Final IgG1 titers at the end of study (day 63) are shown here in comparison with fOVA + PLGA-R. Data were normalized by dividing the titers by naïve mean and are shown here. All data represent the mean ± SEM. All statistical analyses were performed on raw data using one-way ANOVA with Tukey's post-test (**** $P \leq 0.0001$; *** $P \leq 0.001$; ** $P \leq 0.01$ and *ns* indicates not significant).

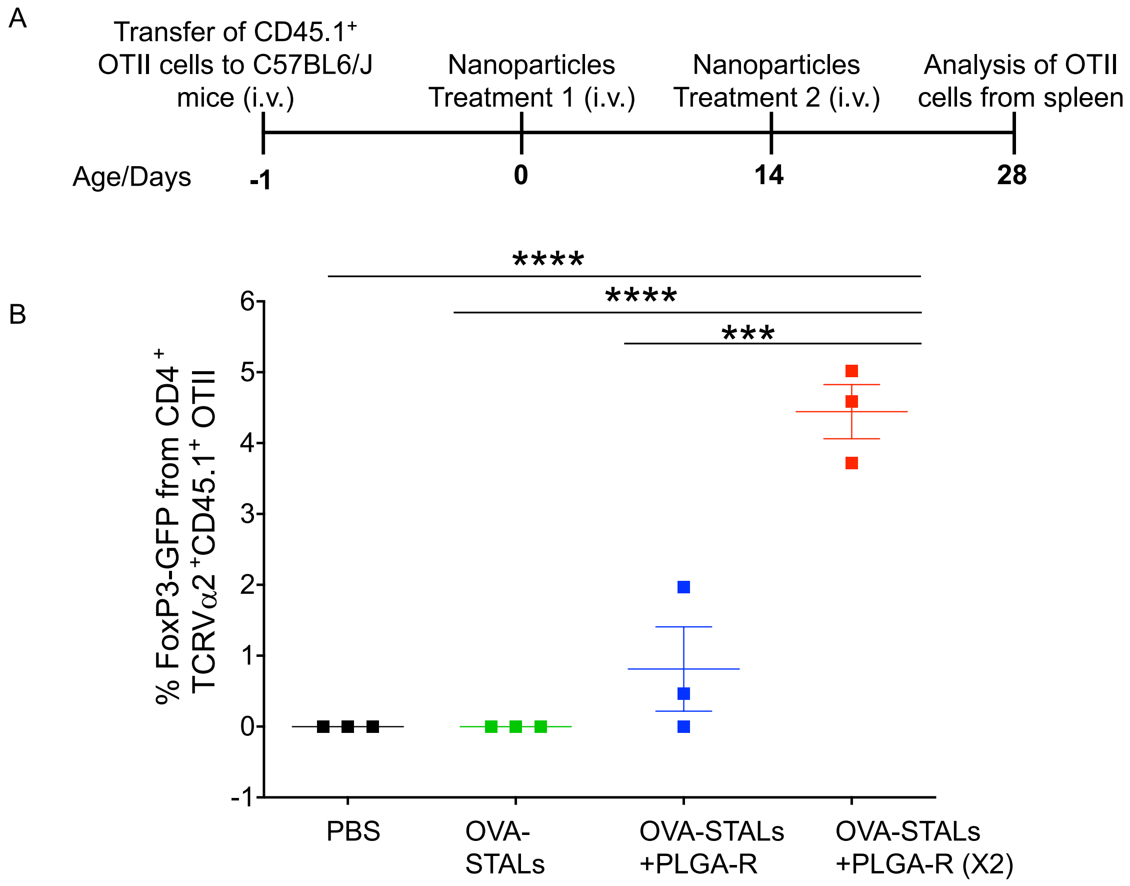


**Figure S2**. Co-delivery of OVA-STALs (OVA-LP-CD22L) and PLGA-R nanoparticles induces regulatory T cells (Tregs). A. Sorted $CD4^{+}FoxP3GFP^{-}CD44^{-}CD62L^{+}$ OTII T cells (0.2 x $10^{6}$) were transferred into $CD45.2^{+}$ C57BL/6J mice on day –1. At day 0, mice (n = 3) received i.v. immunizations with PBS, OVA-STALs alone and OVA-STALs + PLGA-R. Few mice (n = 3) received a second dose of OVA-STALs + PLGA-R on day 14. PLGA-R nanoparticles containing 100 µg of rapamycin. At day 28, spleens were harvested and analyzed by flow cytometry. B. Total percentage of $Foxp3^{+}CD25^{+}$ OTII T cells ($PI^{-}CD19^{-}CD4^{+}TCRV\alpha2^{+}CD45.1^{+}FoxP3GFP^{+}CD25^{+}$) in spleen. All data represent the mean ± SEM. All statistical analyses were performed using one-way ANOVA with Tukey's post-test (**** $p \leq 0.0001$; *** $p \leq 0.001$).

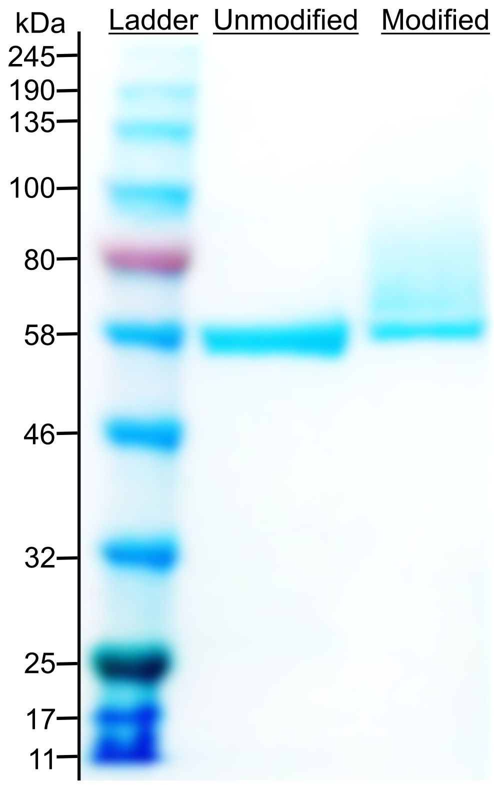


**Figure S3**. SDS-PAGE analysis of GPI protein linked to pegylated lipid. Gel is loaded as follows; lane 1: protein ladder, lane 2: unmodified protein, lane 3: modified protein. Modified protein had between 1-3 lipids attached per protein.
